# Caspase-2 protects against ferroptotic cell death

**DOI:** 10.1101/2023.08.22.553791

**Authors:** Swati Dawar, Mariana C. Benitez, Yoon Lim, Toby A. Dite, Jumana M. Yousef, Niko Thio, Sylvain Garciaz, Thomas D. Jackson, Julia V. Milne, Laura F. Dagley, Wayne A. Phillips, Sharad Kumar, Nicholas J. Clemons

**Author notes:** **Corresponding authors: Nicholas Clemons**, Division of Cancer Research, 305 Grattan Street, Parkville, Victoria 3010, Australia. **E-mail:****; Swati Dawar**, Division of Cancer Research, 305 Grattan Street, Parkville, Victoria 3010, Australia. **E-mail:**. **Conflict of interest:** The authors declare no potential conflicts of interest.

## Abstract

Caspase-2, one of the most evolutionarily conserved member of the caspase family, is an important regulator of the cellular response to oxidative stress. Given that ferroptosis is suppressed by antioxidant defense pathways, such as that involving selenoenzyme glutathione peroxidase 4 (GPX4), we hypothesised that caspase-2 may play a role in regulating ferroptosis. This study provides the first demonstration of an important and unprecedented function of caspase-2 in protecting cancer cells from undergoing ferroptotic cell death. Specifically, we show that depletion of caspase-2 leads to downregulation of stress response genes including *SESN2, HMOX1, SLC7A11* and sensitises mutant-p53 cancer cells to cell death induced by various ferroptosis inducing compounds. Importantly, the canonical catalytic activity of caspase-2 is not required for its role and suggests that caspase-2 regulates ferroptosis via non-proteolytic interaction with other proteins. Using an unbiased BioID proteomics screen, we identified novel caspase-2 interacting proteins (including heat shock proteins and co-chaperones) that regulate cellular responses to stress. Finally, we demonstrate that caspase-2 limits chaperone mediated autophagic degradation of GPX4 to promote survival of mutant-p53 cancer cells. In conclusion, we document a novel role for caspase-2 as a negative regulator of ferroptosis in cells with mutant-p53. Our results provide evidence for a novel function of caspase-2 functions in cell death regulation and open potential new avenues to exploit ferroptosis in cancer therapy.

## Introduction

Caspase-2 is the most conserved member of the caspase family and is well studied for its role as a tumor suppressor under conditions of oncogenic or replicative stress. Previous *in vivo* studies have established that caspase-2 deficient (*Casp2^−/−^*) mice do not develop spontaneous tumors (*1*) but have enhanced susceptibility to tumorigenesis promoted by activation of oncogenes (*K-Ras, EμMyc* and *MMTV/c-neu)* (*2–5*) or loss of a critical tumor suppressor gene *(ATM)* (*6*). Furthermore, studies from human cancers have reported that the *CASP2* gene located on Ch7q is frequently deleted in haematological malignancies (*7*). Moreover, somatic loss-of-function mutations in the human *CASP2* gene, although rare, have been found in multiple cancers (*8, 9*). Mechanistically, the tumor suppressor function of caspase-2 has been attributed to its ability to cause apoptotic removal of aneuploid cells upon replicative stress (*10*). However, absence of caspase-2 is not sufficient to promote tumorigenesis in *MYCN* driven neuroblastoma model (*11*) or following 3-methylcholanthrene (3-MC)-induced fibrosarcoma and irradiation-driven lymphoma (*12*). Together these findings indicate a context specific role of caspase-2 in tumorigenesis.

In addition, *Casp2^−/−^* mice display increased oxidative stress and impaired antioxidant defence response with ageing (*13–15*). Consequently, when challenged with oxidative stressors, *Casp2^−/−^* mice show enhanced oxidative stress induced damage and tumor development (*14*). These findings implicate caspase-2 as an important regulator of the cellular response to oxidative stress. However, in cancer cells, oxidative stress can be a double-edged sword, as excessive accumulation leads to overwhelming damage and ultimately cell death (*16, 17*). This led us to explore the role of caspase-2 in ferroptosis, a non-apoptotic metabolic cell death that occurs from lethal accumulation of iron and lipid peroxides (*18*). Ferroptosis is suppressed by antioxidant defense pathways, such as that involving the selenoenzyme, GPX4 (*18*). Hence, we hypothesised that caspase-2 may play a role in regulating ferroptosis. To investigate this, we used mutant-p53 (mut-p53) cancer cells previously demonstrated to be sensitive to ferroptosis induction due to mut-p53 mediated deregulation of oxidative stress response pathways (*17*). More importantly, mutations in p53 account for more than half of all human cancers, which frequently results in aggressive tumors and poor patient survival (*19*). Therefore, there is an urgent and critical need to develop innovative treatment strategies for patients with mut-p53 cancers.

Here, we have uncovered an important and unprecedented function of caspase-2 in protecting mut-p53 cancer cells from undergoing ferroptotic cell death. Specifically, we found shown that depletion of caspase-2 exquisitely sensitises mut-p53 cancer cells to cell death induced by various ferroptosis inducing drugs. We demonstrate that the catalytic activity of caspase-2 is not required for this role and suggest that caspase-2 regulates ferroptosis via protein-protein interactions, rather than substrate cleavage. We identified novel caspase-2 interacting proteins (including cell chaperone machinery) potentially regulating ferroptosis and we showed for the first time that caspase-2 limits chaperone mediated autophagy (CMA) to protect mut-p53 cancer cells from ferroptosis. We propose inhibiting caspase-2 or its interactions with regulators of ferroptosis as a unique strategy to kill mut-p53 cancer cells by ferroptosis. This provides a paradigm shift in understanding the functions of caspase-2 in cell death and potentially overcomes the limitations of conventional chemotherapy drugs that predominantly act by initiating apoptosis.

## Results

### Acute ablation of caspase-2 enhances ferroptotic cell death in a mut-p53 dependent manner

To examine if caspase-2 plays a role in ferroptosis, we used small interfering RNA (siRNA) to knockdown caspase-2 in isogenic parental p53^null^ and mut-p53 overexpressing (p53^R273H^) lung cancer cells (H1299) and a mut-p53 oesophageal cancer cell line (Flo-1) (Figure 1A). We then measured dose responses to ferroptosis inducing drugs (*17, 18, 20*) erastin, sulfasalazine (SAS), RSL3 and buthionine sulfoximine (BSO) (Supplementary Figure S1A-D). In accordance with our previous studies (*17*), mut-p53 cancer cells were more sensitive to ferroptosis inducers compared to p53^null^ cancer cells (Figure 1B-D). However, both mut-p53 and p53^null^ cancer cells were resistant to doses of BSO up to 5 mM (Fig 1E). Interestingly, we found that depletion of caspase-2 led to enhanced ferroptosis in all cell lines; however, this effect was much more pronounced in the cancer cells expressing mut-p53 (Figure 1B-E). The addition of ferroptosis inhibitors (*18*) deferoxamine (DFO, iron chelator), ferrostatin-1 (Fer1, inhibits lipid peroxidation), N-acetylcysteine (NAC, a cell-permeable analog of cysteine), glutathione-monoethyl ester (GSH-MEE, a cell-permeable analog of GSH), trolox (inhibits lipid peroxidation) and β-mercaptoethanol (reducing agent) rescued erastin mediated cell death following caspase-2 depletion (Figure 1F), confirming that cells were dying from ferroptosis. In contrast, an inhibitor of apoptosis (QVD) had little impact on cell death (Figure 1F), consistent with previous studies (*18*). To further determine if the increased cell death from acute loss of caspase-2 is specific to ferroptosis, we treated mut-p53 and p53^null^ cells with chemotherapeutic drugs 5FU and cisplatin, which cause DNA damage and apoptosis. We found that caspase-2 depletion did not enhance cell death in response to chemotherapeutic drugs (Supplementary Figure S1E-F). Furthermore, knockdown of caspase-3 in mut-p53 cancer cells did not enhance ferroptosis (Supplementary Figure S1 G, H), suggesting that this effect is specific to caspase-2 and is not due to other caspases. Together these results clearly demonstrate that caspase-2 negatively regulates ferroptosis, particularly in mut-p53 cancer cells.

**Figure 1.**
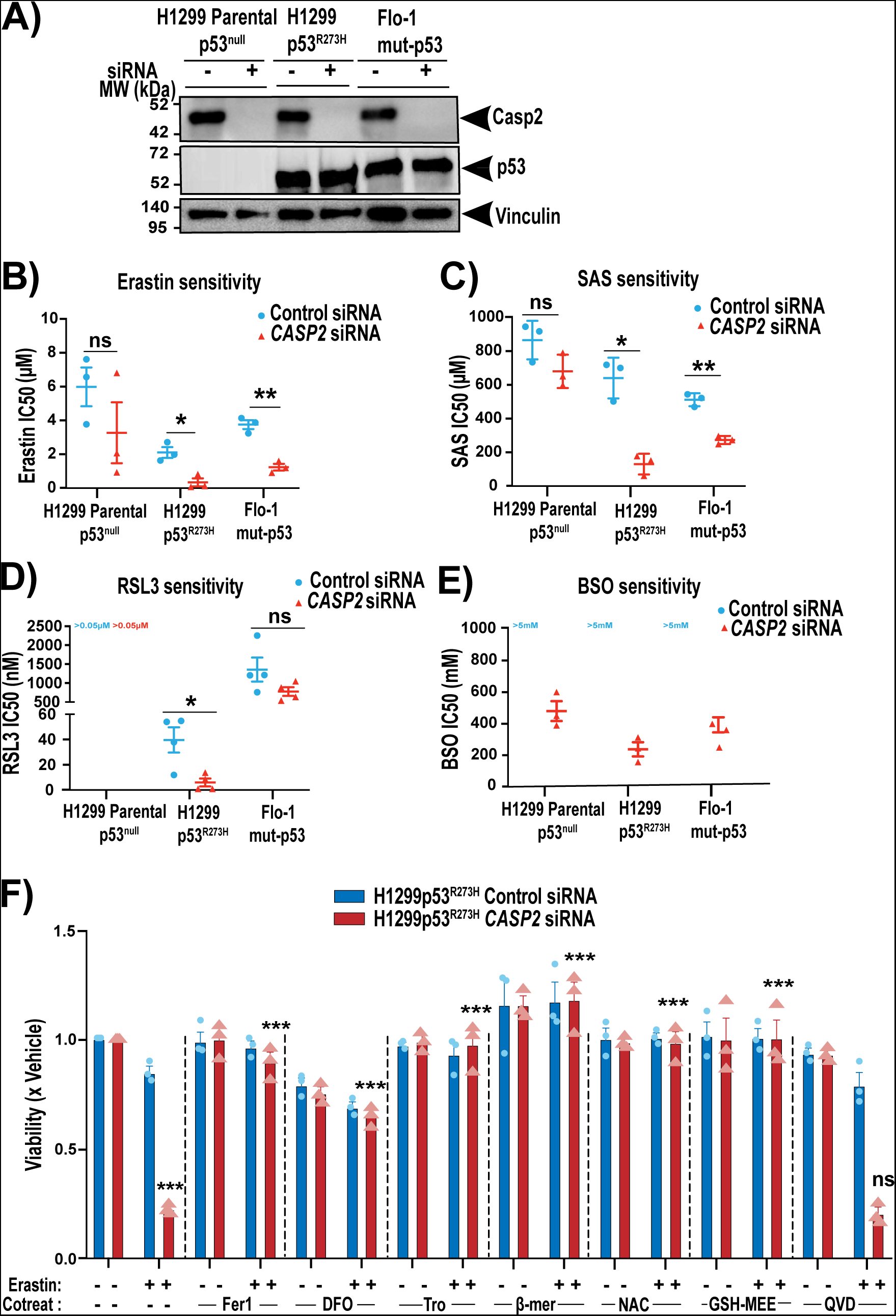
Acute silencing of caspase-2 enhances ferroptotic cell death in a mut-p53 dependent manner and can be rescued by ferroptosis inhibitors. Immunoblot analysis of caspase-2 and p53 expression in H1299 Parental p53^null^, H1299 p53^R273H^ and mut-p53 Flo-1 cells 48 h after transfection with control or *CASP2* siRNA. Vinculin is shown as the loading control. Mean IC50 values after 72 h treatment with (B) erastin, (C) SAS, (D) RSL3 and (E) BSO in H1299 Parental p53^null^, H1299p53^R273H^ and mut-p53 Flo-1 with control or *CASP2* siRNA. (F) Viability of H1299p53^R273H^ cells with control and *CASP2* siRNA at 72 h post treatment with 1 μM erastin, alone or in combination with ferrostatin-1 (Fer-1, 20 μM), deferoxamine (DFO, 100 μM), Trolox (Tro, 1mM), β-mercaptoethanol (β-mer, 100 μM), N-acetyl cysteine (NAC, 5mM), glutathione-methylethyl ester (GSH-MEE, 5mM) or QVD (25 μM). (B-F) Data represented as mean ± s.e.m. from three or four independent experiments. Unpaired *t*-test and one-way ANOVA with Dunnett’s post hoc test were used to estimate significant differences in B-E and F respectively. In F, comparisons were made between erastin only v cotreatment with ferroptosis inhibitors group in *CASP2* siRNA cells. p values are indicated with ns (not significant), \**P* < 0.05, \*\**P* < 0.01, \*\*\**P* < 0.001.

### Loss of caspase-2 leads to a similar increase in ferroptosis in mut-p53 cancer cells

To investigate this phenomenon further we generated H1299p53^R273H^ and Flo-1 cell lines deficient for caspase-2 (H1299p53^R273H^*-CASP2^-/-^*and Flo-1*CASP2^-/-^*) using CRISPR/Cas9 (Figure 2A and Supplementary Figure S2A). Similar to acute silencing of caspase-2 using siRNA, caspase-2 knockout significantly increased the proportion of cells that underwent ferroptosis in response to erastin and RSL3 (Figure 2B, C and Supplementary Figure S2B-E). Furthermore, in the absence of caspase-2, cells formed significantly fewer colonies in comparison to controls in long-term clonogenic assays following treatment with erastin (Figure 2D). In response to ferroptosis inducers, caspase-2 deficient cells showed a ‘ballooning’ phenotype (i.e. formation of a round cell with no cytosol) followed by catastrophic bursting of the plasma membrane, characteristic of ferroptotic cell death (Movies 1-4). Whilst basal levels were unaffected, H1299p53^R273H^-*CASP2^-/-^* cells showed heightened lipid peroxidation (Figure 2E) and reduced total glutathione (Figure 2F) compared to the Cas9 cells following treatment with RSL3 or erastin, respectively, consistent with an enhanced ferroptosis phenotype.

**Figure 2.**
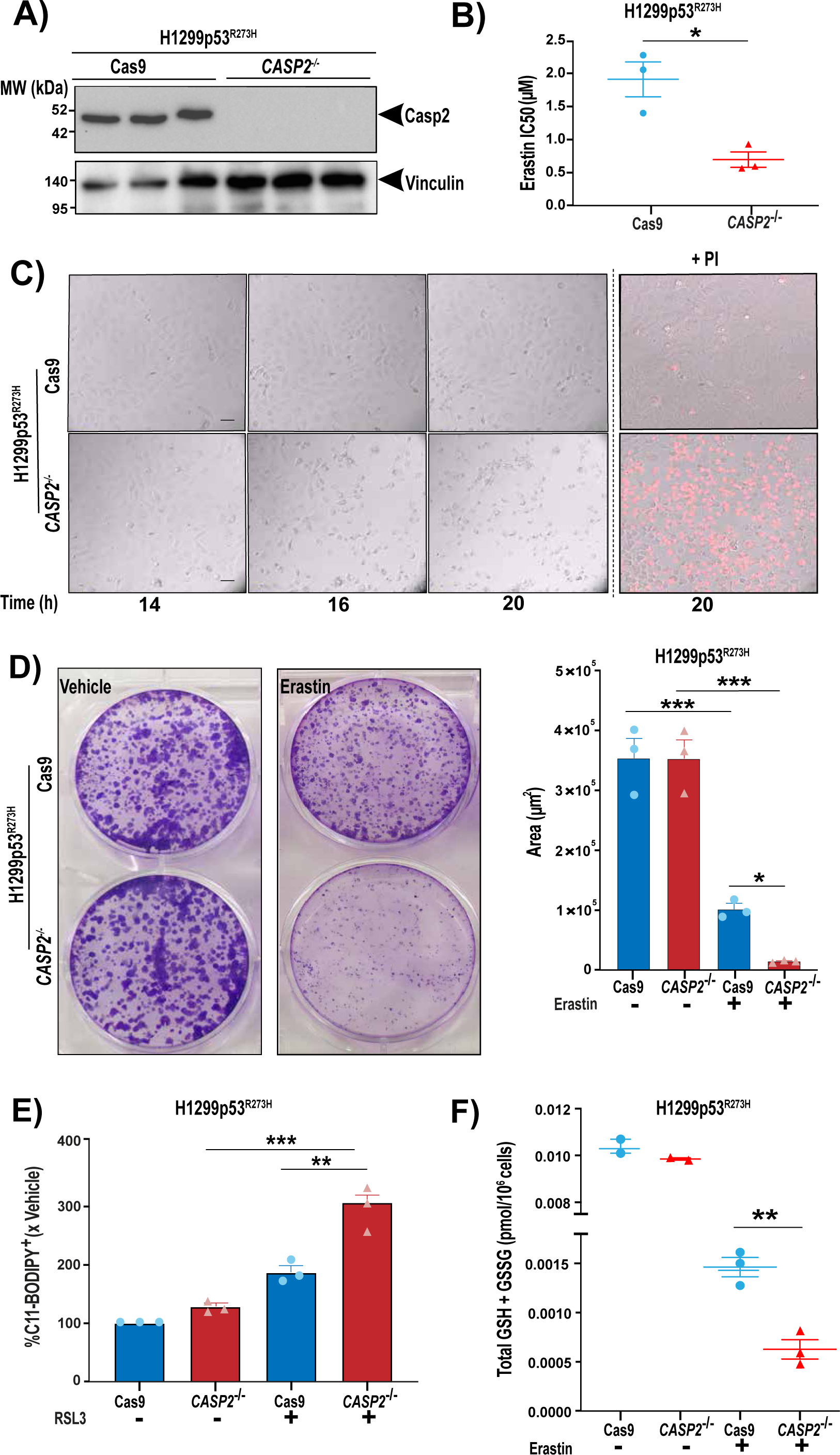
Loss of caspase-2 leads to a similar increase in ferroptosis in mut-p53 cancer cells. (A) Immunoblot analysis of caspase-2 expression in H1299p53^R273H^ cells following CRISPR/Cas9 editing of *CASP2 (CASP2^-/-^)* vs. control cells (Cas9). Vinculin is shown as the loading control. (B) Mean IC50 values (μM) in H1299p53^R273H^ Cas9 and H1299p53^R273H^ - *CASP2^-/-^* cells after 72 h treatment with erastin. (C) Representative bright field images from live cell imaging at the indicated time points of Cas9 and *CASP2*^-/-^ H1299p53^R273H^ cells treated with erastin (2 μM). Right hand panels display dead cells stained red with propidium iodide (PI+). Scale bar=50 μm. (D) Representative images of crystal violet stained colonies of Cas9 or *CASP2*^-/-^ H1299p53^R273H^ cells treated with erastin or vehicle for 12 h, reseeded and cultured over 10 days (*left*). Quantitation of crystal violet stained cell colonies using Cell Profiler software and represented as the area covered by the colonies (*right*). (E) Lipid peroxidation analysis by flow cytometry using C11-BODIPY post RSL3 (40 nM) or vehicle treatment for 18 h in H1299p53^R273H^ Cas9 and H1299p53^R273H^ -*CASP2^-/-^* cells. (F) Total intracellular glutathione (GSH + GSS pmol/10^6^ cells), as determined by GR re-cycling assay after 12 h erastin (2 μM) treatment in H1299p53^R273H^ Cas9 and H1299p53^R273H^ -*CASP2^-/-^* cells. (B, D *right*, E, F) Data represented as mean ±s.e.m. from three independent experiments. Unpaired *t*-test and one-way ANOVA with Tukey’s post hoc test was used to estimate significant differences in B, F and D, E respectively. p values are indicated with \**P* < 0.05 and \*\**P* < 0.01 and \*\*\**P* < 0.001.

To examine the impact of caspase-2 loss in normal p53 wild-type cells, primary mouse embryonic fibroblasts (MEFs) from wild-type and *Casp2^-/-^* mice were treated with erastin and RSL3 (Supplementary Figure S2G). Ferroptotic cell death was not enhanced in *Casp2^-/-^* MEFs compared to wild-type MEFs (Supplementary Figure S2H), thereby supporting the notion that increased sensitivity to ferroptosis upon caspase-2 depletion is specific for mut-p53 cancer cells.

### Loss of caspase-2 alters expression of genes involved in the regulation of cellular stress pathways

Caspase-2 has previously been shown to influence transcriptome changes driven by oncogene activation and tumor suppressor loss in a context specific manner (*4*). To identify the transcriptomic differences and gene enrichment pathways affected by *CASP2* deficiency in mut-p53 cancer cells, we carried out RNA sequencing in H1299p53^R273H^ cancer cells with acute loss and knockout of caspase-2 (Figure 3A, B and Supplementary Figure S3A, B). Comparisons were performed between: (i) control siRNA vs. *CASP2* siRNA, and (ii) Cas9 vs. *CASP2^-/-^* (Supplementary Figure S3A, B). In total, 798 differentially expressed genes (DEGs) with nominal p value < 0.05 were identified in *CASP2* siRNA*/*control siRNA group, of which 157 had a log_2_fold change (FC) of >1 or <-1 (63 upregulated and 94 downregulated). Interestingly, *SLC7A11,* a key component of the glutamate/cystine exchanger (system xCT) and a ferroptosis regulator (*17,18*) was the top downregulated gene in *CASP2* siRNA vs. control siRNA group. In *CASP2^-/-^*/Cas9 group, 1003 significant DEGs were identified with 49 upregulated (log_2_FC >1) and 57 downregulated genes (log_2_FC <-1). There were 67 significant DEGs common in both the groups without caspase-2 as illustrated in the Venn diagram and of these 34 DEGs were either up or downregulated in both models (Figure 3C). These included 14 upregulated (log_2_FC >1) - *ADAM17, APPL2, C12orf11, C5orf15, HMOX1, KIAA1539, MIS12, MKRN1, MIS12, NEU1, RPS7, SESN2, STARD3NL, TSPYL5, ZNF708* and 20 downregulated (log_2_FC <-1) DEGs -*AC008567.1, C20orf27, CD151, CIB1, CPT1A, ECM1, EFCAB4A, FN1, GIPC1, GSTK1, HCFC1, LETMD1, LYD3, MGAT5, MYEOV, PSMB10, SERTAD4, TCF7L1, TIMP2, TOX2* as illustrated in the heat-map (Figure 3D).

**Figure 3.**
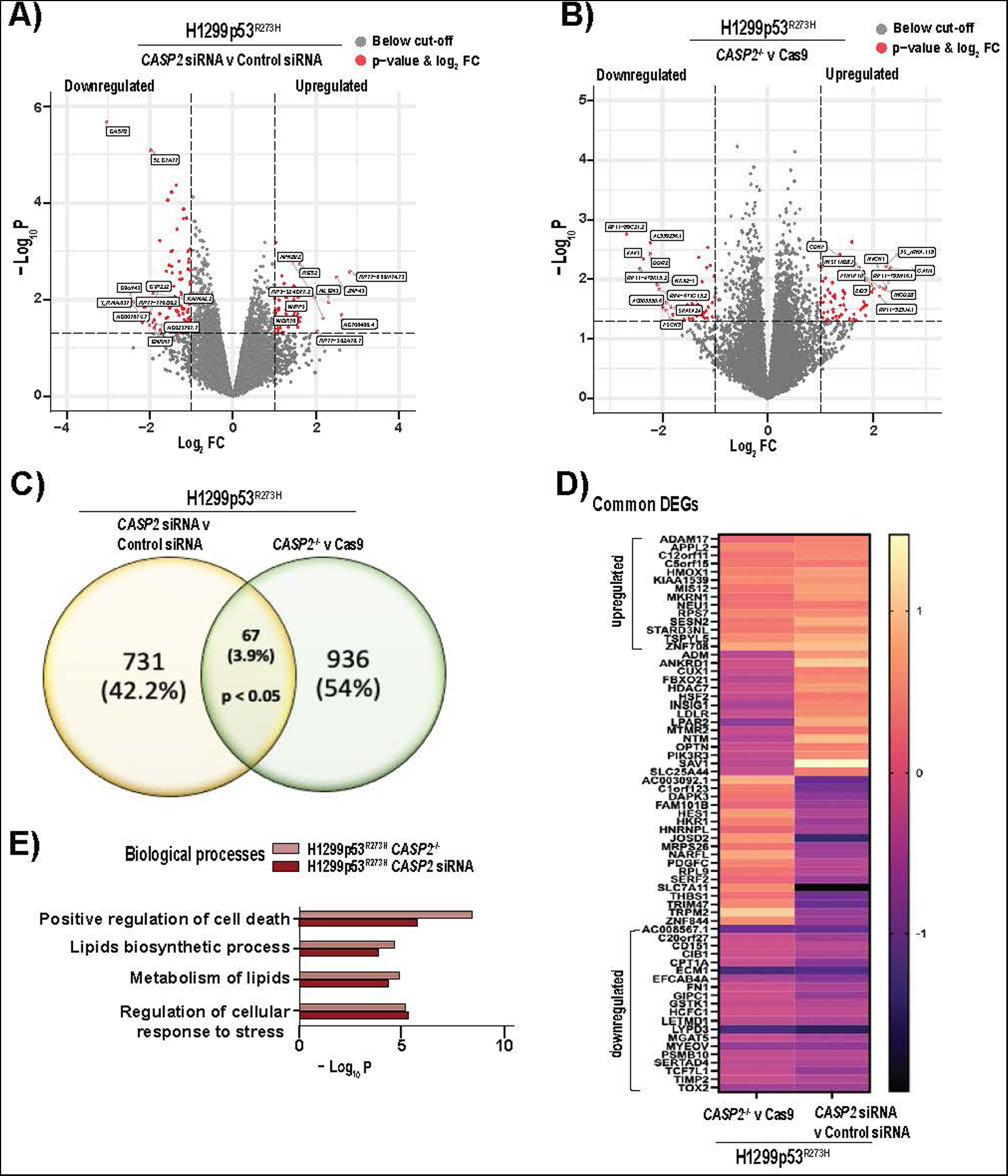
Loss of caspase-2 alters expression of genes involved in the regulation of cellular stress pathways. Volcano plots illustrating DEGs in (A) H1299p53^R273H^ cells transfected with *CASP2* siRNA vs. control siRNA and (B) *CASP2^−/−^* vs. Cas9 H1299p53^R273H^ cells. Red dots indicate DEGs with nominal p value < 0.05 and log_2_FC > or < -1. Top 10 (based on log_2_FC) upregulated and downregulated DEGs are labelled. (C) Venn diagram of unique and overlapping DEGs (p value < 0.05) in *CASP2* siRNA vs. control siRNA group and *CASP2^−/−^* vs. Cas9 group. (D) Heat-map displaying common up/downregulated genes with log_2_FC > 1 or < -1. (E) GO term analysis of the significantly enriched biological processes (nominal p value < 0.01) in common DEGs in the absence of caspase-2 using Metascape (8).

Next, we identified biological processes that were significantly (nominal p value < 0.01) enriched in the DEGs from *CASP2*/control siRNA and *CASP2^-/-^*/Cas9 group. Interestingly, analysis of Gene Ontology (GO) and Reactome terms showed enrichment of biological processes associated with ferroptosis including regulation of cellular responses to stress (GO:0080135), metabolism of lipids (R-HSA-556833), lipids biosynthetic process (GO:0008610), and positive regulation of cell death (GO:0010942) (Figure 3E). Additionally, Gene Set Enrichment Analysis (GSEA) identified downregulation of cellular oxidant detoxification (Supplementary Figure S3C). Together, these data indicate that downregulation of caspase-2 in mut-p53 cancer cells affects expression of genes modulating cellular sensitivity to ferroptosis and therefore, provides a conducive environment for the induction of ferroptotic cell death.

### The catalytic activity of caspase-2 is not required to execute its function in protecting mut-p53 cancer cells against ferroptosis

We wanted to examine if the canonical catalytic activity of caspase-2, which is required for the efficient apoptotic removal of tumorigenic cells, is also required for its role in limiting ferroptosis. To assess this, we re-expressed GFP-tagged caspase-2-C320G (Casp2^C320G^), caspase-2-D135A (Casp2^D135A^) or caspase-2-D330A (Casp2^D330A^) mutants in *CASP2*^-/-^ mut-p53 cancer cells (Figure 4A, B, D). Previous studies have shown that mutation of its catalytic cysteine residue (C320) or aspartate residues (D135 or D330) results in the full (C320 and D135) or partial (D330) loss of its apoptotic activity (*21*). Re-expression of catalytically inactive caspase-2-C320G mutant, or either of the two aspartate mutants, reversed the increased sensitivity to erastin caused by knockout of endogenous wild-type caspase-2 (Figure 4C, E), clearly demonstrating that the catalytic activity and apoptotic function of caspase-2 is not required for its role in preventing ferroptosis. We, therefore, hypothesised that caspase-2 regulates ferroptosis via non-proteolytic interaction with other proteins.

**Figure 4.**
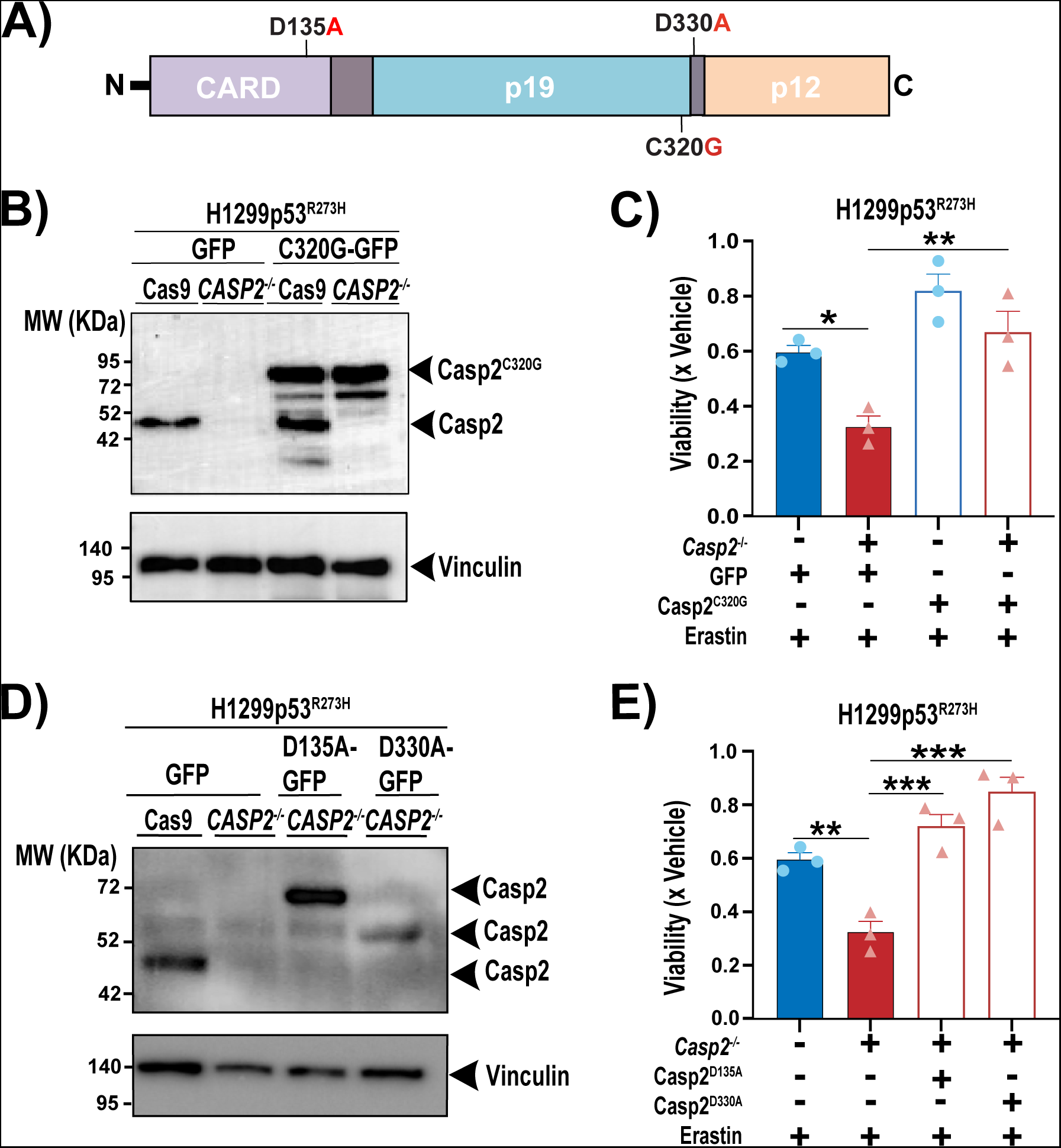
The catalytic activity of caspase-2 is not required to execute its function in protecting mut-p53 cancer cells against ferroptosis. (A) Diagram of caspase-2 consisting of prodomain or caspase recruitment domain (CARD), long subunit (p19) and small subunit (p12) along with the specific mutation sites introduced. (B) Immunoblot analysis of caspase-2 in H1299p53^R273H^ Cas9 and H1299p53^R273H^-*CASP2^-/-^* cells transfected with a GFP-tagged catalytically inactive caspase-2-C320G (Casp2^C320G^/ C320G-GFP) expression plasmid or control plasmid (GFP). The higher MW of the ectopic Casp2^C320G^ is because of the GFP tag. Vinculin is shown as the loading control. (C) Viability in H1299p53^R273H^ Cas9 and H1299p53^R273H^-*CASP2^-/-^* cells ectopically expressing Casp2^C320G^ or GFP 24 h post treatment with erastin (2 μM) normalized to vehicle treated cells. (D) Immunoblot analysis of caspase-2 in H1299p53^R273H^ Cas9 cells transfected with GFP control plasmid, and H1299p53^R273H^-*CASP2^-/-^* cells transfected with GFP-tagged caspase-2-D135A (Casp2^D135A^/D135A-GFP) and caspase-2-D330A (Casp2^D330A^/D330A-GFP) expression plasmids or GFP control plasmid. Vinculin is shown as the loading control. The 60-kDa band in Casp2^D330A^ mutant cells represents a partially processed form of caspase-2-GFP protein (5, 6). (E) Viability in H1299p53^R273H^ Cas9 cells and H1299p53^R273H^-*CASP2^-/-^* cells ectopically expressing Casp2^D135A^ and Casp2^D330A^ at 24 h post treatment with erastin (2 μM) normalized to vehicle treated cells. (C, E) Data represented as mean ±s.e.m. from three independent experiments. One-way ANOVA with Bonferroni’s post hoc test was used to estimate significant differences in (C, E). p values are indicated with \**P* < 0.05, \*\**P* < 0.01 and \*\*\**P* < 0.001.

### Identification of novel caspase-2 interacting proteins underpinning ferroptotic cell death in mut-p53 cancer cells

To identify proteins that interact with caspase-2 that could potentially explain its ability to regulate ferroptosis, we performed an unbiased BioID proteomics screen. An *E. coli*-derived promiscuous biotin ligase (BirA*; BirA-R118G mutant) was fused with the Casp2^C320G^ mutant as a bait to biotinylate proteins in close proximity. We did not use wild-type caspase-2 since it leads to apoptosis when expressed ectopically (*21*). A BirA* only expression vector was used as a control to validate the specificity of caspase-2 interacting proteins. Re-expression of BirA*-Casp2^C320G^ mutant in H1299p53^R273H^-*CASP2^-/-^* cells was confirmed by immunoblotting (Supplementary Figure S4A). In line with our previous results, re-expression of catalytic inactive caspase-2-BirA* mutant partially rescued erastin mediated ferroptotic cell death in H1299p53^R273H^-*CASP2^-/-^* cells (Supplementary Figure S4B).

BioID was performed in the presence and absence of erastin to capture basal interacting proteins and determine whether the caspase-2 interactome was altered in response to erastin. Biotinylated proteins were captured from protein lysates using streptavidin agarose beads and depletion of biotinylated proteins in the supernatant was confirmed by immunoblotting (Supplementary Figure S4C, D). Potential caspase-2-BirA* interacting proteins bound to the beads were isolated and identified by mass spectrometry-based proteomic analysis. There were >250 proteins identified within each sample (Supplementary Figure S4E, Supplementary table 1). A protein was determined to have significant differential expression if the log_2_FC was ≥ 1 and the adjusted p value was ≤ 0.05. As expected, caspase-2 was the most enriched protein under both basal and erastin treated conditions. Excluding caspase-2, we identified 25 caspase-2-interacting candidate proteins under basal conditions (Figure 5A). Following treatment with erastin, we identified 17 enriched caspase-2 interacting candidate proteins, of which 14 were part of the basal interactome, and 3 additional proteins namely KRT77, CTAG1A and SQSTM1 (Figure 5B).

**Figure 5.**
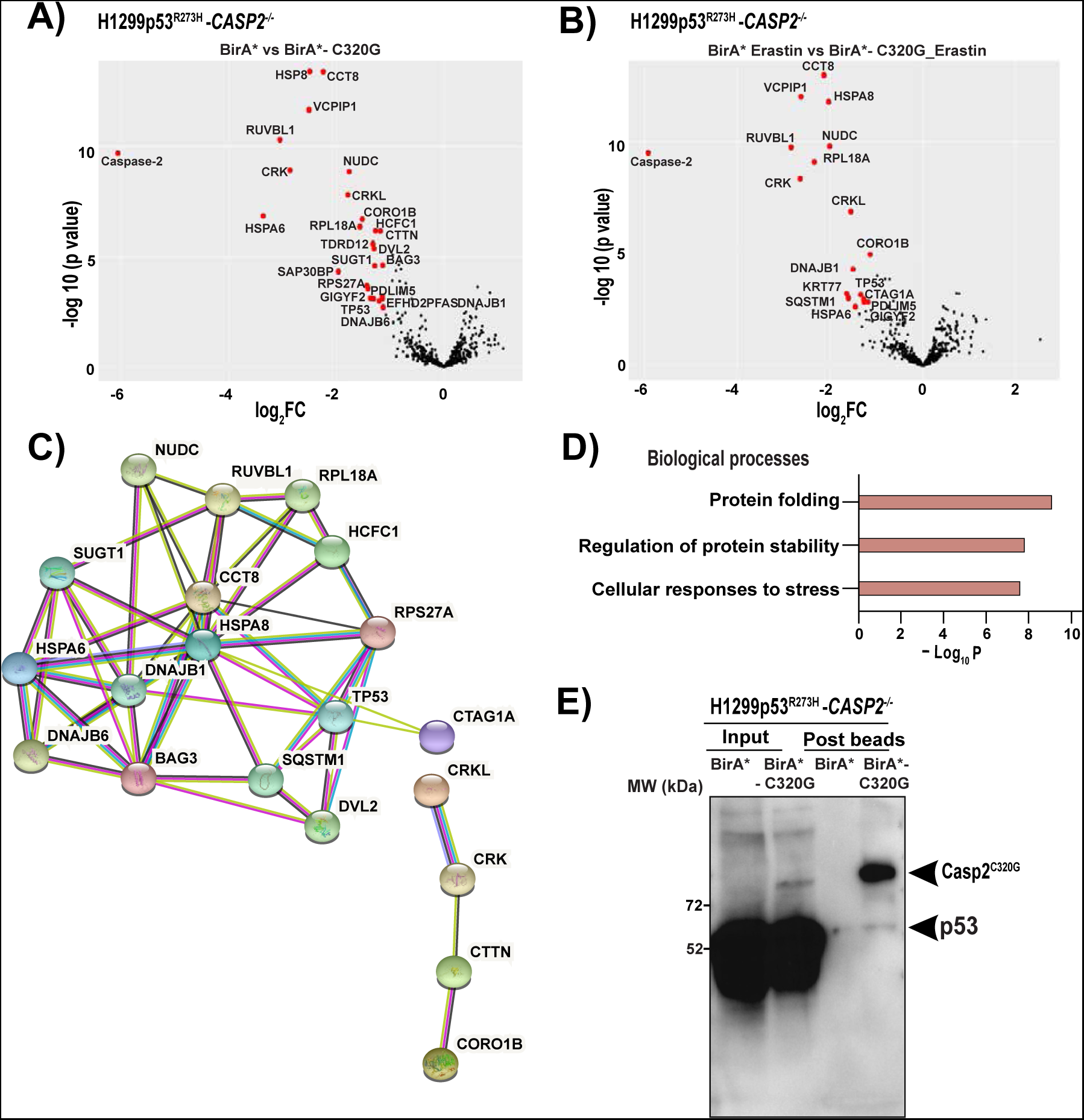
Identification of novel caspase-2 interacting proteins underpinning ferroptotic cell death in mut-p53 cancer cells. Volcano plot illustrating the log_2_ fold changes of biotinylated proteins for BirA* control vs. BirA*-Casp2^C320G^ for (A) untreated samples and (B) erastin treated samples. Five biological replicates per group were prepared for MS analysis. Proteins were deemed to exhibit differential expression if the log2 fold change in protein expression was ≥ 1-fold and an adjusted p value ≤ 0.05. (C) Protein-protein interaction network among top 28 significant caspase-2 interacting proteins, based on STRING annotations. Edges indicate a range of protein-protein associations (physical and functional) such as experimentally determined (pink), curated databases (cyan), text mining (light green), co-expression (black), and protein homology (purple). Disconnected nodes (n=8 proteins) in the network are not shown. (D) Functional enrichment for biological processes (p value < 0.01) associated with caspase-2 interacting proteins as determined by Metascape. (E) Co-immunoprecipitation and immunoblotting for mut-p53 in H1299p53^R273H^-*CASP2^-/-^* cells expressing BirA* control and BirA*-Casp2^C320G^ before and after pull-down with streptavidin agarose beads under basal conditions.

To classify the 28 unique caspase-2 interacting proteins into significant interaction networks we utilized STRING database with a medium confidence score of > 0.4, which returned 52 biological connections (enrichment p value < 0.001) depicted as edges between the nodes (Figure 5C). To further classify the resulting network into functional groups, we performed functional enrichment for biological processes. Interestingly, we identified caspase-2 interacting proteins to function in processes related to protein folding and stability (GO: 0006457, GO: 0031647) and cellular responses to stress (R-HSA-3371556) (Figure 5D). This included Hsp70 chaperones (HSPA8, HSPA6) and a suite of Hsp70/Hsp90 co-chaperones (DNAJB1, BAG3, DANJB6, SUGT1, NUDC and RUVBL1), suggesting that caspase-2 might protect mut-p53 cancer cells from ferroptosis through interaction with chaperone machinery.

Interestingly, our quantitative proteomics data identified mut-p53 as a caspase-2 interacting protein. Extending from this, we validated the interaction by immunoblotting and confirmed that mut-p53 immunoprecipitated with caspase-2 in BirA*-Casp2^C320G^ mutant expressing H1299p53^R273H^-*CASP2^-/-^* cells (Figure 5E). Therefore, our data demonstrated for the first time a direct interaction between caspase-2 and mut-p53 under basal growth conditions, which may explain the mut-p53 selective modulation of ferroptosis by caspase-2 we observed (Figure 1).

### Caspase-2 limits chaperone mediated autophagic degradation of GPX4

The major cognate chaperone, Hsc70/HSP73 (HSPA8), is the core regulator of chaperone mediated autophagy (CMA), a form of selective autophagy that also requires Hsp90 (*22*). CMA has previously been shown to be associated with ferroptosis execution via degradation of GPX4 (*23*). Therefore, we next investigated if loss of caspase-2 promotes ferroptosis via chaperone mediated autophagic degradation of GPX4. Interestingly, we observed reduced GPX4 protein levels in cells with acute knockdown of caspase-2 compared to control cells (Figure 6A). Moreover, treatment with 17AAG, an inhibitor of HSP90 chaperone activity, could reverse the increased sensitivity to erastin treatment in *CASP2* siRNA H1299p53^R273H^ cells (Figure 6B). Furthermore, treatment with 17AAG partially reversed caspase-2 knockdown mediated downregulation of GPX4 protein levels (Figure 6C). We next determined whether knockdown of caspase-2 can also regulate autophagy in general. Our results did not show an overall increase in autophagy in caspase-2 depleted cells compared to control cells as detected by light chain 3 (LC3)B-II and SQSTM1 protein levels before and after treatment with bafilomycin A1 (Baf A1), an inhibitor of autophagic flux (Supplementary Figure S5). Taken together, these findings suggest that loss of caspase-2 promotes chaperone mediated autophagic degradation of GPX4 which provides a sensitized environment and when challenged with ferroptosis inducing drugs leads to severe lipid peroxidation and cell death in mut-p53 cancer cells (Figure 6D).

**Figure 6.**
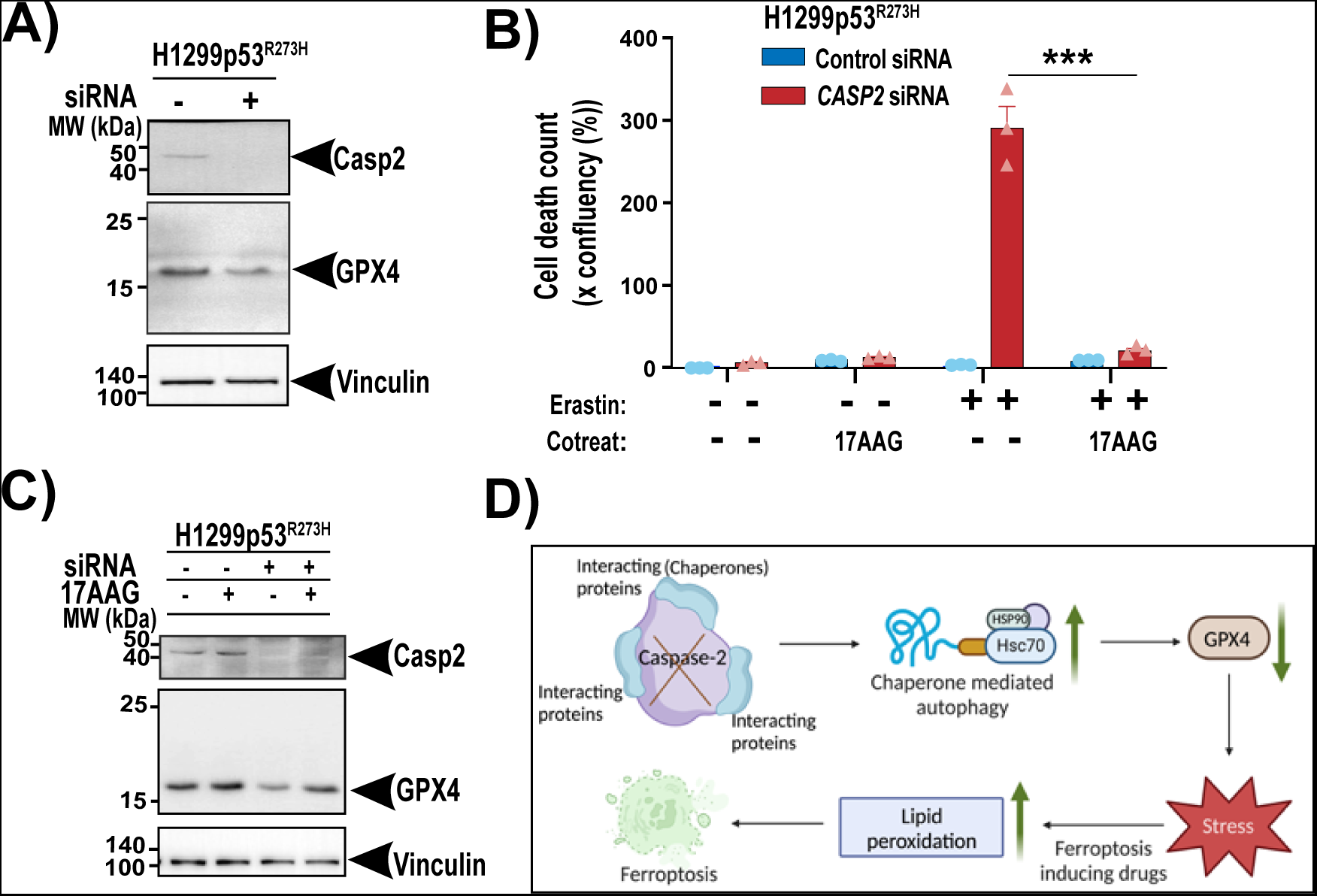
Caspase limits chaperone mediated autophagic degradation of GPX4. (A) Immunoblot analysis of GPX4 and caspase-2 expression in H1299p53^R273H^ control or *CASP2* siRNA cells. Vinculin is shown as the loading control. (B) Viability of H1299p53^R273H^ cells with control and *CASP2* siRNA at 18 h post treatment with 1 μM erastin, alone or in combination with 17AAG (500 nM). (C) Immunoblot analysis of GPX4 and caspase-2 expression in H1299p53^R273H^ control or *CASP2* siRNA cells following treatment with 17AAG. Vinculin is shown as the loading control. (D) Schematic diagram summarizing the role of caspase-2 in ferroptosis. Data represented as mean ± s.e.m. from three independent experiments. Unpaired *t*-test was used to estimate significant differences in B. p value is indicated with \*\*\**P* < 0.001.

## Discussion

In this study, we provide the first demonstration of a previously unrecognized function of caspase-2 in protecting cancer cells carrying mut-p53 from ferroptosis. Our findings show that regulation of ferroptosis by caspase-2 does not require its canonical catalytic activity but instead involves protein-protein interactions. Moreover, we identified components of core chaperone machinery as key caspase-2 interacting proteins and provide evidence that caspase-2 protects cells from ferroptosis at least partially via preventing chaperone-mediated autophagic degradation of GPX4. We propose that targeting caspase-2 creates a previously unknown Achilles’ heel that could potentially be exploited as a treatment to selectively kill mut-p53 cancer cells by ferroptosis.

Several studies have previously implicated caspase-2 in non-apoptotic contexts, including its role as an important regulator of cellular oxidative stress pathways (*14, 24, 25*). A predominant feature of *Casp2^-/-^* mice is enhanced oxidative damage to proteins, lipids and reduced antioxidant signaling with ageing, making *caspase-2* deficient mice more susceptible to these stress conditions (*13–15*). In view of these findings, we explored the possibility of caspase-2 in regulating ferroptosis, a non-apoptotic cell death known to occur in response to imbalance in oxidative stress and antioxidant defense pathways (*18, 26*). Our cell viability and imaging data indicate that loss of caspase-2 renders mut-p53 cancer cells more susceptible to pro-ferroptotic drugs resulting from reduced GSH levels and enhanced lipid peroxidation. The RNA sequencing data identified differential expression of oxidative stress related genes including *SESN2, HMOX1, SLC7A11* in mut-p53 cancer cells without caspase-2 under basal conditions. These observations are consistent with previous studies implicating association of caspase-2 loss with transcriptional regulation of genes involved in tumorigenesis and redox signaling (*4, 14*). Collectively, these studies imply that caspase-2 loss supports a favourable environment for ferroptosis and does not simply act by enhancing kinetics of erastin effect.

Previous studies have reported a context specific role for caspase-2 in tumorigenesis. Although several studies employing mouse models of tumorigenesis support a tumor suppressor role for caspase-2 (*2, 3, 5, 6, 8, 9, 27*), cancer promoting role for caspase-2 has been reported for *MYCN* driven neuroblastoma in a mouse model (*11*). Consistent with this study, our data describe an oncogenic role for caspase-2 in limiting ferroptosis in mut-p53 cancer cells. The mechanism for caspase-2 tumor suppressor function has been mostly attributed to its ability in preventing cells from becoming cancerous through activating apoptosis or cell cycle arrest through MDM2 cleavage and p53 stabilisation (*28*). Both these roles are dependent on the catalytic activity of caspase-2 (*10, 29*). Furthermore, as a proteolytic enzyme, caspase-2 is known to be inactive in dividing cells and activated via dimerization following a variety of cellular stress stimuli to mediate cleavage of cellular substrates to drive apoptosis (*10, 30–33*). Our novel finding suggests contrasting roles for caspase-2 in ferroptosis and apoptosis and that dimerization and proteolytic activity of caspase-2 is not required for the regulation of ferroptosis.

Studies so far have entirely focused on identifying substrates cleaved by caspase-2 in apoptosis, whereas investigations to identify targets and regulators of caspase-2 in non-apoptotic contexts are limited. We successfully utilised BioID proximity-dependent biotinylation to identify caspase-2 interacting proteins in mut-p53 cancer cells. Our interactome data revealed novel binding partners of caspase-2 including heat shock chaperone proteins, co-chaperones and p53 (in our case mut-p53) under basal and erastin treated conditions. Caspase-2 has been previously reported to be activated by PIDDosome-like complex comprising of PIDD1 (p53-inducible with a death domain1) and RAIDD (RIP-associated ICH-1/CED-3 homologous proteins with a death domain) in apoptosis (*34*). Activated caspase-2 then cleaves MDM2 to stabilise p53 or BID to generate tBID, leading to cell cycle arrest or apoptosis (*28, 35*). In contrast, for the first time, our results reveal caspase-2 and mut-p53 as binding partners in regulating ferroptosis. Our cell viability data further supports this finding as caspase-2 loss particularly sensitises mut-p53 cancer cells and to a less extent p53 null cancer cells or wild-type p53 cells. Further studies are required to validate this interaction in other cancer cells with mutated p53.

CMA promotes degradation of oxidatively damaged and functional proteins to maintain cellular homeostasis (*23*). A recent study reported that activation of CMA led to GPX4 degradation during ferroptosis (*23*). In line with this finding, we found that caspase-2 limits CMA degradation of GPX4 to promote survival of mut-p53 cancer cells. In contrast, there was no evidence to suggest a change in autophagic flux/protein degradation via bulk autophagy. A role of caspase-2 as a negative regulator of autophagy under basal conditions has been previously shown (*25*). It was demonstrated that loss of caspase-2 increased autophagy as a protective mechanism to promote cell survival against oxidative stress (*25*). In contrast, we show that it could be a vulnerability if the stress promotes ferroptosis. Specifically, we demonstrate that loss of caspase-2 provides a sensitized environment conducive for ferroptosis that triggers severe oxidative stress to mediate cell death. Interestingly, SQSTM1, a cargo receptor on autophagosomes that is involved in a wide range of “sub-types” of autophagy (*36*), was one of the interacting candidates in our Bio-ID screen. However, SQSTM1 has not been implicated in CMA. Whether loss of caspase-2 inhibits other selective types of autophagy associated with ferroptosis (such as alters levels of iron, ferritinophagy), and also if mut-p53 is having a role in determining the context of autophagy, requires further investigation.

Despite being the most evolutionary conserved of caspases, the physiological function of caspase-2 has been enigmatic (*37, 38*). This study provides a whole new perspective in the unanticipated role of caspase-2 as a negative regulator of ferroptosis, independent of its proteolytic activity. Overall, our findings provide a proof-of-concept study for targeting caspase-2 and/or its interacting proteins as a promising novel therapeutic strategy to promote killing of mut-p53 cancer cells by ferroptosis.

## Materials and Methods

### Drugs and general chemicals

Erastin, RSL3, BSO, 17AAG and Baf A1 were from SelleckChem (Houston, Texas, USA). Cisplatin and 5FU were from Hospira (Lake Forrest, Illinois, USA). GSH-MEE was from Cayman Chemicals (Ann Arbor, Michigan, USA). GSH, β-mercaptoethanol, NAC, trolox, ferrostatin-1, QVD, DFO and SAS were from Sigma-Aldrich (St. Louis, Missouri, USA). Unless otherwise specified, all general chemicals were from Sigma-Aldrich.

### Cell cultures

Isogenic lung cancer (H1299p53^null^ and H1299p53^R273H^) and esophageal cancer cell lines (Flo-1) were maintained in a humidified incubator at 37 °C with 5% CO_2_ in RPMI 1640 medium from Life Technologies (Carlsbad, California, USA) containing 2.5 mM _L-_glutamine and high-glucose DMEM (Life Technologies), respectively, supplemented with 10% heat inactivated fetal bovine serum (FBS) (Sigma-Aldrich) and 100 U ml^-1^ Penicillin/100 μg ml^-1^ Streptomycin (Life Technologies).

Primary MEFs were derived from wild-type and *Casp2*^−/−^ embryos at day 13.5 as previously described (*10, 27, 39*) and maintained in high-glucose DMEM with 0.2 mM L-glutamine (Sigma-Aldrich), 10% FBS (JRH Biosciences, Lenexa, Kansas, USA) supplemented with 100 μM penicillin/streptomycin (Sigma-Aldrich). All the cell lines tested negatively for mycoplasma infection and their identities confirmed by STR analysis.

### siRNA transfections and dose response assays

Cells were transfected with 40 nM *Casp2* (ON-TARGET plus Human CASP2 siRNA SMARTpool L-003465-00-0050), *Casp3* (ON-TARGET plus Human CASP3 siRNA SMARTpool L-004307-00-0020) or non-targeting control siRNA pools (ON-TARGET plus non-targeting pool D-001810-10-05) using Lipofectamine RNAiMAX transfection reagent (Life Technologies) as per the manufacturer’s instructions. Knockdown was confirmed by immunoblotting.

For dose response assays in 384-well microplates (Falcon^®^), a BioTek EL406 personal liquid handling workstation was used for dispensing cells and a Caliper Sciclone ALH3000 robot dispensed the siRNA:lipid:opti-MEM complexes. In brief, 500 cells per well were reverse transfected on day 1 for 48 h incubation. On day 3, a 10-point log_2_ serial dilution of drugs were added to the cells for a further 72 h incubation. For cell viability measurements, 4 μl of AlamarBlue^®^ reagent (Life Technologies) was added per well, incubated for 4 h and fluorescence was read at 550 nm/590 nm on a Cytation 3 Imaging Reader (BioTek, Winooski, Vermont, USA).

For ferroptotic death rescue, 3500 cells per well in 100 µl culture media were seeded in 96-well plates (Falcon^®^) and reverse transfected with Control and *CASP2* siRNA for 48 h at 37 °C. The culture medium was then replaced with media containing 1 µM erastin and/or ferroptosis inhibitors including 20 µM ferrostatin-1, 1 mM trolox, 100 µM β-mercaptoethanol, 25 µM QVD, 5 mM NAC, 5 mM GSH-MEE, or 100 µM DFO followed by incubation for 72 h at 37 °C and cell viability was assessed with AlamarBlue^®^ as above.

For CMA rescue, 3500 cells per well in 100 µl culture media were seeded in 96-well plates (Falcon^®^) and reverse transfected with Control and *CASP2* siRNA for 48 h at 37 °C. The culture medium was then replaced with media containing 1 µM erastin and/or CMA inhibitor 17AAG followed by incubation for 48 h at 37 °C and cell viability was assessed with Incucyte FLR (Essen BioScience). Percentage (%) of cell death was determined by dividing the percentage of red confluency by the percentage of phase confluency.

### Cell viability assay in MEF lines

For cell viability assay, two independently generated MEF lines per genotype at passage 2-4 were used. Briefly, cells were seeded in triplicate at 3000 cells per well in 50 µl complete MEF culture media in a 96-well microplate (BD Biosciences, Franklin Lakes, New jersey, USA) and cultured overnight at 37 °C with 10% CO_2_. Next day culture medium (50 µl) containing DMSO, 1 µM erastin or 400 nM RSL3 was added, followed by 12 h incubation at 37 °C with 10% CO_2_. Twenty-five microliters of MTS/PMS (96:4) reagent was added to each well, followed by incubation for 3 h in a 10% CO_2_ incubator at 37 °C. Absorbance was measured at 490 nm using a FLUOstar Omega (BMG Labtech, Ortenberg, Germany). Control wells (culture media only) were used to detect the cell-free background absorbance.

### Generation of *CASP2* knockout cell lines using CRISPR/Cas9 and dose response assays

*CASP2* knockout cell lines in H1299p53^R273H^ and Flo-1 were generated using Alt-R CRISPR-Cas9 technology (Integrated DNA Technologies, Coralville, Iowa, USA) as per manufacturer’s instructions. Briefly, cells were transduced to express constitutive Cas9 endonuclease (FUCas9Cherry, Victorian Centre for Functional Genomics, Peter Mac, Australia) and mCherry positive cells were sorted by flow cytometry. *Casp2* crRNA (Hs.Cas9.CASP2.1.AA/AltR1/rGrCrUrGrUrUrUrGrGrCrUrArGrCrArCrCrArCrUrGrUrUrU rUrArGrArGrCrUrArUrGrCrU/AltR2/) and Alt-R^®^ CRISPR-Cas9 tracrRNA duplexes were combined to a final concentration of 1 µM. To generate *CASP2* knockout cell lines, Cas9 expressing H1299p53^R273H^ and Flo-1 were seeded at 2 x 10^5^ cells in 6-well plates (Falcon^®^) and reverse transfected with crRNA:tracrRNA complex using Lipofectamine RNAiMAX solution for 48 h. Absence of caspase-2 was confirmed by immunoblotting.

For cell viability assay in CRISPR Cas9 generated *CASP2* knockout cell lines, cells were seeded at 500 cells per well in 100 µl culture medium in a 384-well microplate and cultured overnight at 37 °C with 5% CO_2_. Next day, media was changed and a 10-point log_2_ serial dilutions of drugs were added to the cells for a further 72 h incubation. Twenty microlitres of AlamarBlue^®^ was added to 100 µl of culture medium, incubated for 4 h and fluorescence was read at 550 nm/590 nm on a Cytation 3 Imaging Reader (BioTek).

### GSH assay

Total intracellular glutathione measurements were performed as previously described (*40*). Briefly, 5 × 10^5^ H1299p53^R273H^ Cas9 and H1299p53^R273H^-*Casp2*^-/-^ cells with and without 2 µM erastin treatment for 12 h were homogenized in ice-cold 10 mM HCl. Proteins were precipitated by adding 5-sulfosalicylic acid (Sigma-Aldrich) to a final concentration of 1%. Samples were centrifuged to remove precipitates and supernatants were collected and stored at -20 °C until analysis. Samples were incubated in the presence of 5,5’-dithio-bis-[2-nitrobenzoic acid] (DTNB, 0.73 mM) (Sigma-Aldrich), EDTA (4 mM) (Sigma-Aldrich), dihydronicotinamide-adenine dinucleotide phosphate (NADPH, 0.24 mM) (Sigma-Aldrich), 110 mM sodium phosphate (NaH_2_PO_4_) buffer (pH 7.4) and glutathione reductase (GR; Sigma-Aldrich) from baker’s yeast (1.2 U/ml). The absorbance at 412 nm was measured every 15 sec for 5 min on a Cytation 3 Imaging Reader (BioTek). Total concentration of GSH + GSSG was calculated based on a GSH standard curve.

### Lipid peroxidation detection assay

Lipid peroxidation was detected using C11-BODIPY(581/591) dye (ThermoFisher Scientific). Briefly, 2 x 10^5^ H1299p53^R273H^ Cas9 and *Casp2*^-/-^ cells were seeded in 6-well plates and incubated with and without RSL3 at 40 nM for 18 h. The cells were washed twice with PBS and incubated with C11-BODIPY (5 µM) for 30 minutes at 37 °C with 5% CO_2_. The cells were collected, washed twice with PBS and resuspended in 200 µL of PBS with propidium iodide (2µg/ml) and analysed using flow cytometry (BD LSRFortessa, BD Biosciences).

### Live cell imaging

H1299p53^R273H^ Cas9 and *Casp2*^-/-^ cells were seeded in 96-well plate at 3500 cells and treated with 2 µM erastin or vehicle (DMSO) and imaged for 18 h using Olympus IX 83 (Tokyo, Japan). Propidium iodide (2µg/ml) was used as an indicator of cell death. Each frame was captured per 10 minutes of real-time.

### Clonogenic survival assay

H1299p53^R273H^ Cas9 and *Casp2*^-/-^ cells at 1 × 10^5^ were seeded in a 35 mm dish and treated with 2 µM erastin or vehicle for 12 h. Cells were then harvested and reseeded at a density of 10^3^ viable cells per well in a 6-well plate. After culturing for 10 days, the colonies were stained with 0.5% crystal violet (Sigma-Aldrich) and area was counted using Cell Profiler software (Tokyo, Japan). A colony was defined as consisting of at least 50 cells.

### Plasmid construction, transfection and viability

pEGFPN1 or pEGFPN1-Casp2^C320G^ constructs have been described previously (*5*). pEGFPN1-Casp2^D330A^, pEGFPN1-Casp2^D135A^ mutant constructs were generated by GenScript. H1299p53^R273H^-*CASP2^-/-^* cells were seeded at 2 x 10^5^/well in 6-well plates one day before transfection. Next day, the cells were transfected with the respective constructs using Fugene^®^ HD from Promega (Madison, Wisconsin, USA) or Lipofectamine 3000 (Life Technologies) transfection reagent as per the manufacturer’s instructions. Cells were incubated with the transfection complexes for 24 h and GFP positive cells were sorted by flow cytometry. Sorted cells were then reseeded in 96-well plates and cell viability with and without erastin was measured by AlamarBlue^®^. Expression of caspase-2 mutants was confirmed by immunoblotting.

### Immunoblotting

Protein lysates were prepared from cells in RIPA buffer (25 mM Tris/HCl pH 7.4, 150 mM NaCl, 1% nonyl-phenoxylpolyethoxylethanol, 1% sodium deoxycholate, 0.1% sodium dodecyl sulphate) in the presence of protease/phosphatase inhibitor cocktail (Thermo Scientific, Rockford, Illinois, USA). Homogenates were further treated by three freeze/thawed cycles in liquid nitrogen, clarified by centrifugation at 13.2K rpm and protein concentration was determined by BCA assay (Bio-Rad, Hercules, California, USA). For western blot analysis, 20-30μg of lysates was resolved on a 4-15% or 10% precast gels (Bio-rad) and transferred onto 0.45μm PVDF membrane (Immobilon-P; Merck Millipore) and probed with the specified antibody overnight at 4°C. Secondary antibodies, conjugated with Horseradish peroxidase (HRP) were incubated at room temperature for 2 h. Proteins were visualized using ECL Plus (Pierce) or Immune-Star Western Chemiluminescence kit (Bio-rad). The following antibodies were used: caspase-2 (clone 11B4) (In-house), GPX4 antibody (Abcam, Cambridge, Massachusetts, USA), caspase-3 (Cell Signalling Technology, Danvers, Massachusetts, USA), p53 (DO-1 and 1801) (In-house), LC3-B (Cell Signalling Technology, Danvers, Massachusetts, USA) and Vinculin (Abcam, Cambridge, Massachusetts, USA).

### RNA-sequencing

Total RNA was isolated from H1299p53^R273H^ Control and *CASP2* siRNA transfected cells, H1299p53^R273H^ Cas9 and *Casp2*^-/-^ cells using the Nucleospin RNA kit (Macherey-Nagel GmbH & Co, Germany). Three biological replicates each were prepared. RNA concentration and purity were measured with a NanoDrop1000^TM^ spectrophotometer (Thermo Fisher Scientific). RNA-seq was performed by the Molecular Genomic core facility at Peter MacCallum Cancer Centre. Library was prepared with QuantaSeq Lexogen 3’mRNAFW on the Illumina NextSeq 500 system using Single End 75 base reads and read depth between 3-4 million reads per sample.

RNA-sequencing reads were aligned using HISAT2 (*41*), and gene expression was quantified using HTSeq software (*42*). Normalised expression was measured in count-per-million (CPM) in log2 scale, with library size adjustment using Trimmed Mean of M-values method (*43*), using edgeR (*43*) R package. Differential expression analysis was performed using LIMMA-Voom workflow (*44*). Genes with consistent extreme-low expression across 80% of samples were excluded from analysis, to reduce noise amplification in Voom method. Gene set enrichment analysis was performed using GSEA (*45*), comparing two groups of triplicates. MSigDB gene set collections were used as reference to discover significantly up-regulated gene sets. Heatmap figures are visualisation of normalised expression data, using Pearson correlation distance to compare across genes, and Euclidean distance to compare across samples. Colour scale of each gene is centered at the average normalised expression across samples in the heatmap.

### Statistical analysis of data

All statistical analyses were carried out using GraphPad Prism, Version 9.2 (GraphPad, GraphPad Software Inc) or Microsoft Excel, Version 16.56. The difference between data groups was analysed using two-tailed Student’s *t*-test or ANOVA with Dunnett’s multiple comparisons post hoc test as indicated in the manuscript. Data are represented as mean ± standard error of the mean (s.e.m.). p values < 0.05 were considered as statistically significant.

### BioID

Point directed mutagenesis was performed in BirA*-Casp2 to generate BirA*-Casp2^C320G^ mutant by GenScript (Piscataway, New Jersey, USA). H1299p53^R273H^-*Casp2*^-/-^ cells were seeded at 1.5 x 10^6^ cells in 10cm dishes one day before transfection. Next day, the cells were transfected with 14 µg of constructs in RPMI without antibiotics and incubated at 37 °C for 30 h. The following day cells were re-seeded at 8 x 10^5^ cells and treated with biotin (500 µM) and erastin at 3 µM for 18 h at 37 °C with 5% CO_2_. Cells were washed eight times with ice-cold PBS and then re-suspended in RIPA buffer with phosphatase and protease inhibitors. Five biological replicates per condition was prepared. Protein was quantified using BCA assay (Bio-rad). Biotinylated proteins were captured from lysates (500 µg per replicate) using 10 μg streptavidin sepharose (GE healthcare, Waukesha, Wisconsin, USA) for 8 h, rotating at 4° C. The beads were then washed three times with RIPA lysis buffer, transferred to Snap Cap Spin Columns (Pierce, Rockford, Illinois, USA) and washed three times with high salt buffer (10 mM Tris-HCl, 500 mM NaCl, pH7.5), then three times with tryptic digest buffer (50 mM ammonium bicarbonate, 1.5 M urea). Proteins were digested on-beads for 16 hours at 37 °C using 1 µg trypsin and peptides collected into new tubes by centrifugation. The collected peptides were lyophilized to dryness using a CentriVap (Labconco, Kansas, Missouri, USA), before reconstituting in 10 µl 0.1% formic acid (FA)/2% acetonitrile (ACN) ready for mass spectrometry analysis. BirA* and BirA*-Casp2^C320G^ mutant expression in H1299p53^R273H^-*Casp2*^-/-^ was confirmed by western blotting and viability following erastin treatment was assessed by AlamarBlue^®^ as mentioned previously. Depletion of biotinylated proteins from lysates was confirmed by immunoblotting using Streptavidin HRP antibody (STAR 5B, Bio-rad).

### Mass spectrometry analysis

Peptides (3 µl) were separated by reverse-phase chromatography on a C_18_ fused silica column (inner diameter 75 µm, OD 360 µm × 15 cm length, 1.6 µm C_18_ beads) packed into an emitter tip (IonOpticks, Fitzroy, Victoria, Australia), using a nano-flow HPLC (M-class, Waters, Milford, Massachusetts, USA) coupled to a timsTOF Pro (Bruker, Billerica, Massachusetts, USA) equipped with a CaptiveSpray source. Peptides were loaded directly onto the column at a constant flow rate of 400 nl/min with buffer A (99.9% Milli-Q water, 0.1% FA) and eluted with a 30-min linear gradient from 2 to 34% buffer B (99.9% ACN, 0.1% FA). The timsTOF Pro was operated in diaPASEF mode using Compass Hystar 5.1. Settings were as follows: Mass Range 100 to 1700m/z, 1/K0 Start 0.6 V·s/cm^2^ End 1.6 V·s/cm^2^, Ramp time 100ms, Lock Duty Cycle to 100%, Capillary Voltage 1400V, Dry Gas 3 l/min, Dry Temp 180 °C. The acquisition scheme for diaPASEF is shown in supplementary table 2. The collision energy was ramped linearly as a function of the mobility from 59 eV at 1/K0 = 1.6 V·s/cm^2^ to 20 eV at 1/K0 = 0.6 V·s/cm^2^. The mass spectrometry proteomics data have been deposited in the ProteomeXchange Consortium via the PRIDE partner repository with the dataset identifier PXD041339. with the username and password K7eRYyXN (*46*).

### Raw data processing and analysis

Data files were analysed by DIA-NN v1.8 software (*47*). Data was searched against the human Uniprot Reference Proteome with isoforms (downloaded August 2021), with recombinant protein sequences added, as a FASTA digest for library-free search with a strict trypsin specificity allowing up to 2 missed cleavages. The peptide length range was set to 7-30 amino acids. Precursor charge range was set between 1-4, and m/z range of 300-1800. Carbamidomethylation of Cys was set as a fixed modification. Precursor FDR was set to 1% and match between runs was on.

Data cleaning and analysis were performed using R (version 4.1.2). Proteins without any proteotypic precursors or with q-value greater than 0.01 were removed. In addition, only proteins that were quantified in at least 50% of replicates in at least one condition were kept. The protein intensities were log_2_-transformed. Missing values were imputed by Missing At Random (v2-MAR) method implemented in msImpute package (v. 1.3.3). The data were normalized using cyclicloess method implemented in limma (v. 3.50.0). Differential analysis was performed using limma (v. 3.50.0). A protein was determined have significant differential expression if the log_2_ FC was ≥ 1 and the adjusted p value was ≤ 0.05. The following comparisons were made: BirA* vs BirA*-Casp2^C320G^ and BirA*_Erastin vs BirA*-Casp2^C320G^_Erastin. The R-package ggplot2 (v. 3.3.5) was used to visualize the results.

## Acknowledgements

We thank all the members of our laboratory for discussions and useful comments. Specifically, we thank Karen Montgomery for help with dose response experiments, Metta Jenna from Centre for Advanced Histology and Microscopy core facility within Peter Mac, for help with live cell imaging. Lastly, we thank GenScript for generating caspase-2 mutant constructs and BirA*-Casp2^C320G^ construct.

## Author contributions

S.D. conceptualised, designed the study, carried out most experiments, collected and analysed data, M.C.B. performed and analysed dose response assays in siRNA transfected cells, Y.L., carried out cell viability assay in MEF lines, contributed BioID reagents and provided technical advice, T.D., J.M.Y, L.F.D., provided technical advice on BioID, carried out mass spectrometry and data analysis, N.T. carried out RNA sequencing data analysis, S.G. helped with performing and analysis of lipid peroxidation detection assay using flow cytometry, T.D.J carried out Immunoblotting and cell viability assay for CMA experiment, J.M. carried out Immunoblotting repeats for CMA experiments, S.K. provided reagents and guidance, W.P. provided constructive feedback and guidance, N.C. supervised the study and interpreted the data, S.D. and N.C. drafted and edited the manuscript. All authors gave comments and approved the final manuscript.

