## Supplementary Figure Legends for "Caspase-2 protects against ferroptotic cell death"

Supplementary figure 1

(A) Viability at 72 h in control or *CASP2* siRNA transfected H1299 parental $p53^{null}$, H1299 $p53^{R273H}$ and mut-p53 Flo-1 cells post treatment with the indicated doses of (A) erastin, (B) SAS, (C) RSL3, (D) BSO, (E) 5FU and (F) cisplatin. (G) Immunoblot analysis of caspase-3 expression in H1299$p53^{R273H}$ after 48 h transfection with control or *CASP2* siRNA. Vinculin is shown as the loading control. (H) Viability in control or *CASP3* siRNA transfected H1299 $p53^{R273H}$ cells at 72 h post treatment with the indicated doses of erastin. (A-F, H) Data represented as mean ±s.e.m. from three or four independent experiments.

Supplementary figure 2

Viability in Cas9 and *CASP2*$^{-/-}$ H1299$p53^{R273H}$ cancer cells at 72 h post treatment with (A) erastin or (B) RSL3. IC50 values (nM) after treatment with RSL3 are shown (B, *right*). (C) Immunoblot of caspase-2 expression in Flo-1 cells following CRISPR/Cas9 editing of *CASP2* *(CASP2*$^{-/-}$*)* vs. control cells (Cas9). Vinculin is shown as the loading control. Viability in Cas9 and *CASP2*$^{-/-}$ Flo-1 cancer cells at 72 h post treatment with (D) erastin or (E) RSL3. IC50 values are shown to the right. (F) Representative flow cytometry plots for lipid peroxidation analysis (C11 BODIPY fluorescence) of Cas9 and *CASP2*$^{-/-}$ H1299$p53^{R273H}$ cells treated with RSL3 (40 nM) for 18 h. (G) DNA agarose gel image of genotyping and Immunoblot analysis of caspase-2 in primary MEFs from WT and *Casp2*$^{-/-}$ mice (two independent MEF lines for each genotype). Vinculin is shown as the loading control (H) Viability at 12 h post erastin (1 μM) and RSL3 (0.4 μM) treatment in WT and *Casp2*$^{-/-}$ MEFs. (A, B, D, E, H) Data represented as mean ±s.e.m. from three or four independent experiments. Unpaired *t*-test was used to estimate significant differences in B and D. One-way ANOVA with Bonferroni's post hoc test was used to estimate significant differences in H. p values are indicated with ns and $^{*}P < 0.05$.

Supplementary figure 3

Heat-maps displaying upregulated (red) and downregulated (blue) DEGs from three independent replicates in (A) *CASP2*$^{-/-}$ vs. Cas9 H1299$p53^{R273H}$ cells and (B) H1299$p53^{R273H}$ cells transfected with *CASP2* siRNA vs. control siRNA. (C) GSEA score curves for the identified upregulated and downregulated pathways in *CASP2* siRNA vs. control siRNA group and *CASP2*$^{-/-}$ vs. Cas9 group. Black bars represent the position of members of the category in the ranked list together with the respective p value.

Supplementary figure 4

(A) Immunoblot analysis of caspase-2 in H1299$p53^{R273H}$-*CASP2*$^{-/-}$ cells transfected with BirA*-$Casp2^{C320G}$ or BirA* control plasmid. Vinculin is shown as the loading control. (B) Viability in H1299$p53^{R273H}$-*CASP2*$^{-/-}$ cells expressing BirA* control or BirA*-$Casp2^{C320G}$ re-expression at 24 h post treatment with erastin (2 μM). Immunoblot analysis of biotinylated proteins in H1299$p53^{R273H}$-*CASP2*$^{-/-}$ cells expressing BirA* control or BirA*-$Casp2^{C320G}$ before and after pull-down with streptavidin agarose beads under (C) basal conditions or (D) after treatment with erastin. Depletion of biotinylated proteins was confirmed by streptavidin antibody. Vinculin is shown as the loading control. (E) Bar graph indicating number of proteins identified per sample for each of the groups. Proteins that were identified in 60% of samples (3 out 5 samples) in at least one group were kept for analysis. Data in (B) is represented as mean±s.e.m. from three independent experiments. Unpaired *t*-test was used to estimate significant differences in (B). p values are indicated with $*P < 0.05$.

Supplementary figure 5

Immunoblot analysis of LC3B-I, LC3B-II, SQSTM1 and caspase-2 expression in H1299 $p53^{R273H}$ control or *CASP2* siRNA cells following treatment with 2 h post treatment with Baf A1 (200 nM). Vinculin is shown as the loading control.

Movie Legends

Movie 1
Live imaging in Cas9 cells treated with erastin and imaged for 18h. Each frame was captured per 10 minutes of real-time. Related to Figure 2.

Movie 2
Live imaging in $CASP2^{-/-}$ H1299$p53^{R273H}$ cells treated with erastin and imaged for 18h. Each frame was captured per 10 minutes of real-time. Related to Figure 2.

Movie 3
Live imaging in Cas9 H1299$p53^{R273H}$ cells treated with erastin and Propidium Iodide. Cells were imaged for 18h and each frame was captured per 10 minutes of real-time. Related to figure 2.

Movie 4
Live imaging in $CASP2^{-/-}$ H1299$p53^{R273H}$ cells treated with erastin and Propidium Iodide. Cells were imaged for 18h and each frame was captured per 10 minutes of real-time. Related to figure 2.
