## Supplementary figures and images for "Caspase-2 protects against ferroptotic cell death"

### Supplementary Figure S1

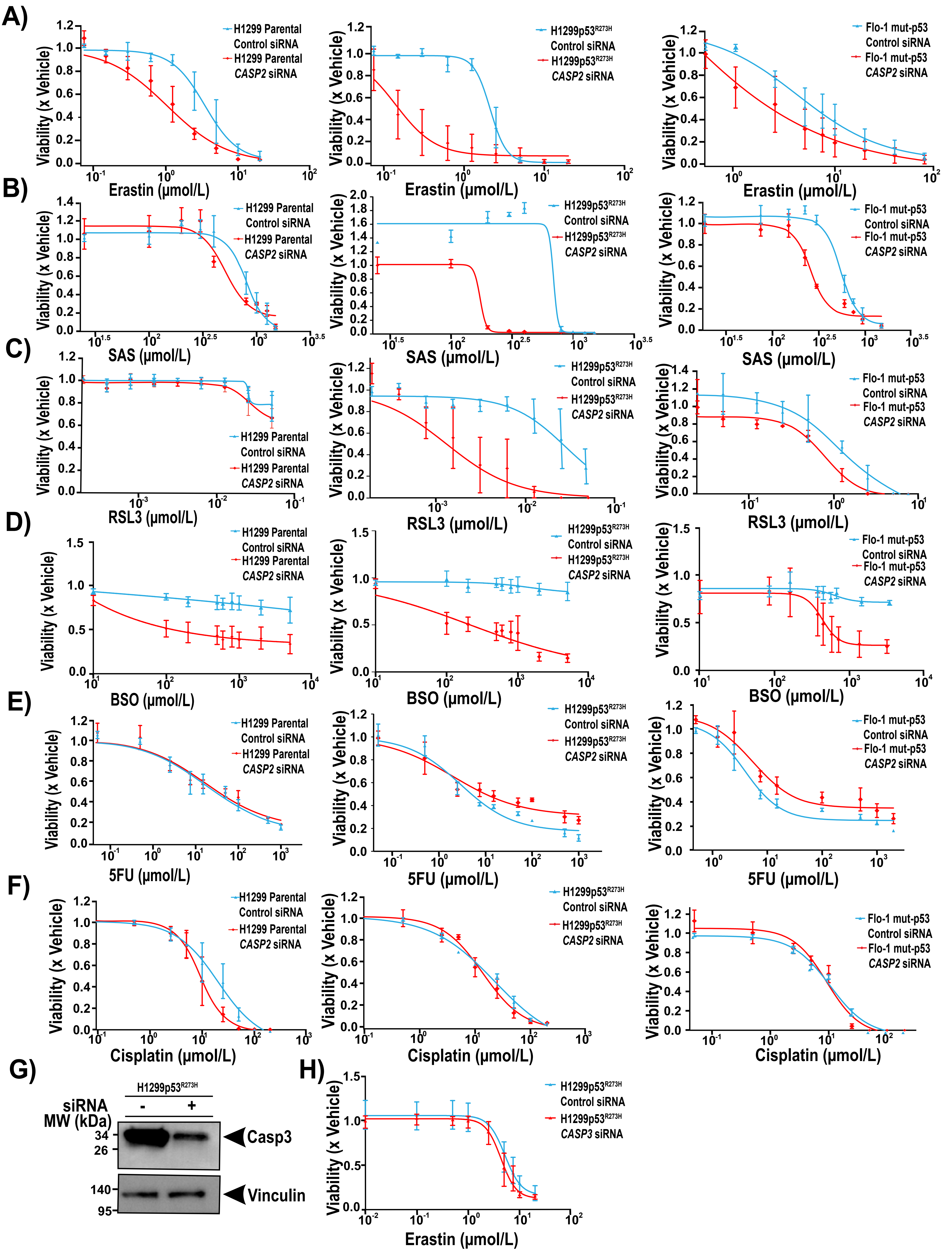

### Supplementary Figure S2

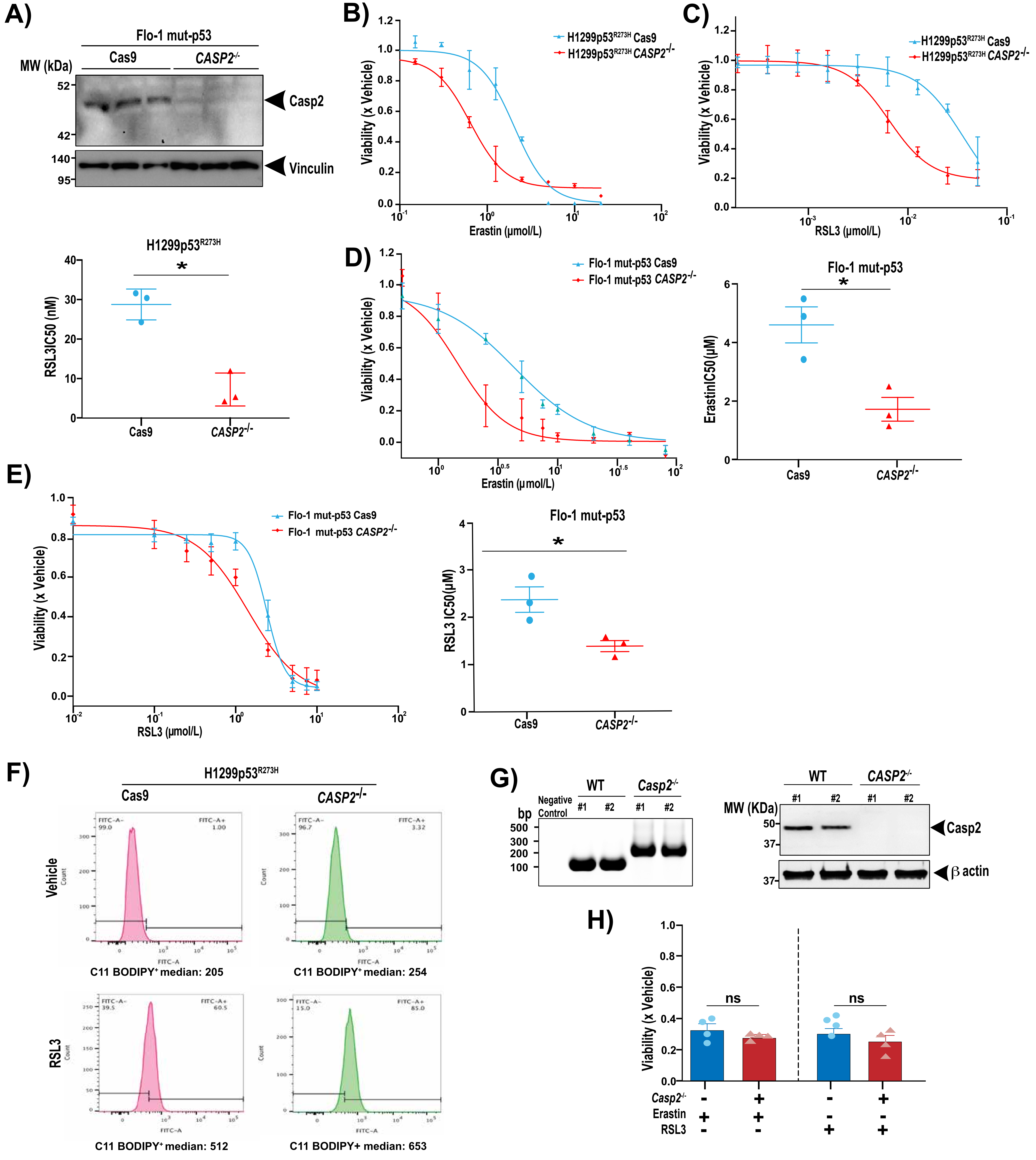

### Supplementary Figure S3

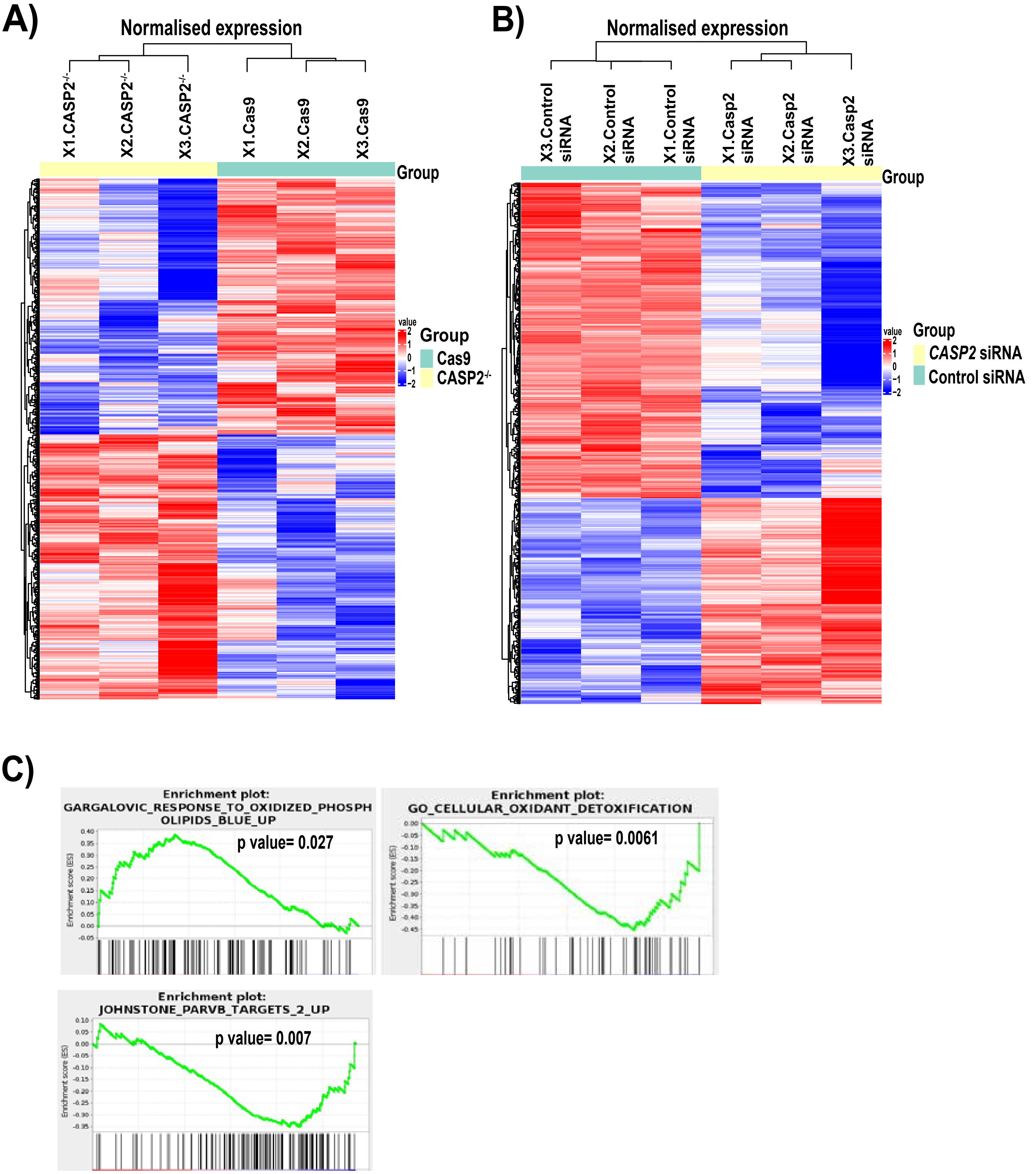

### Supplementary Figure S4

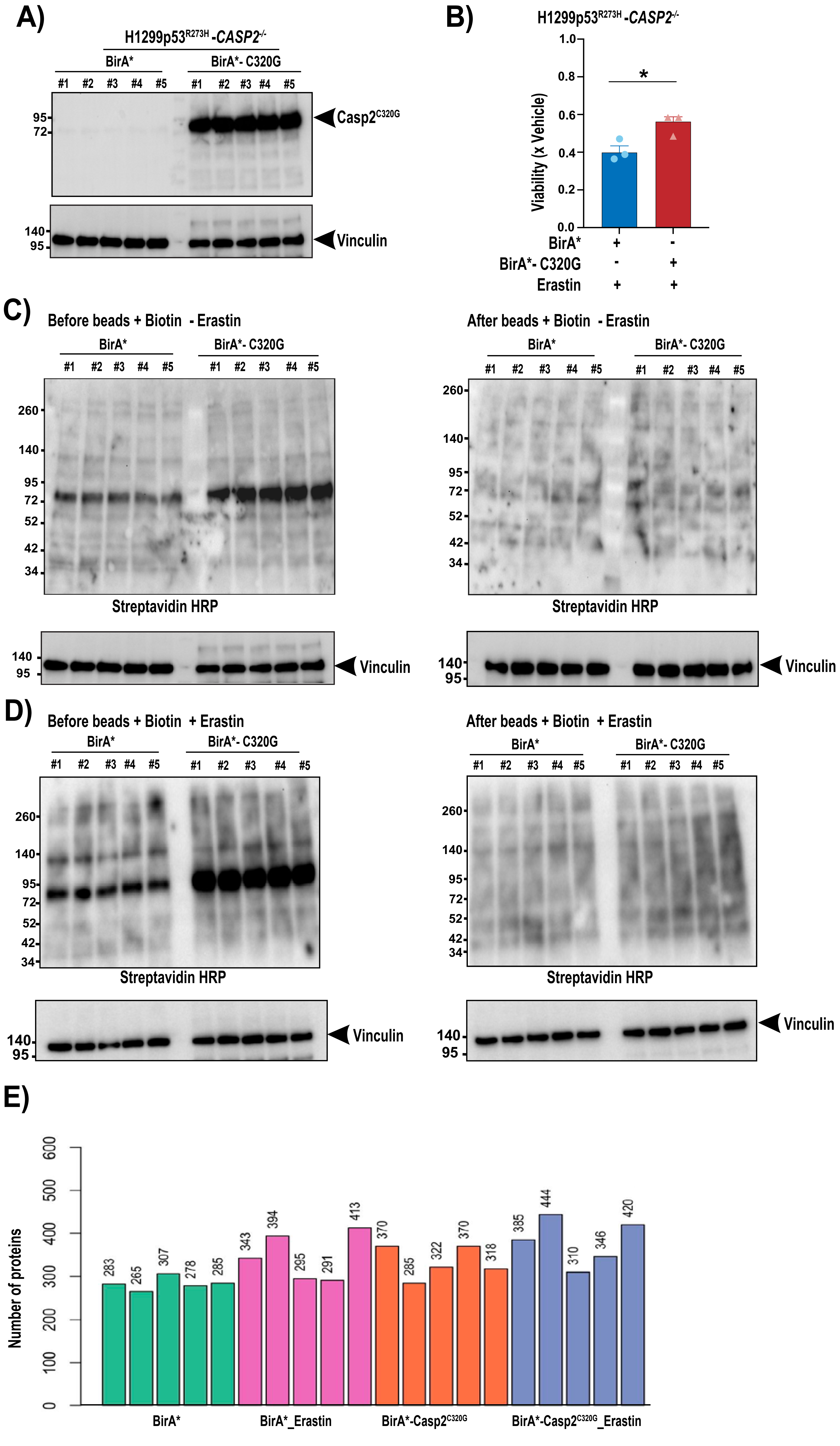

### Supplementary Figure S5

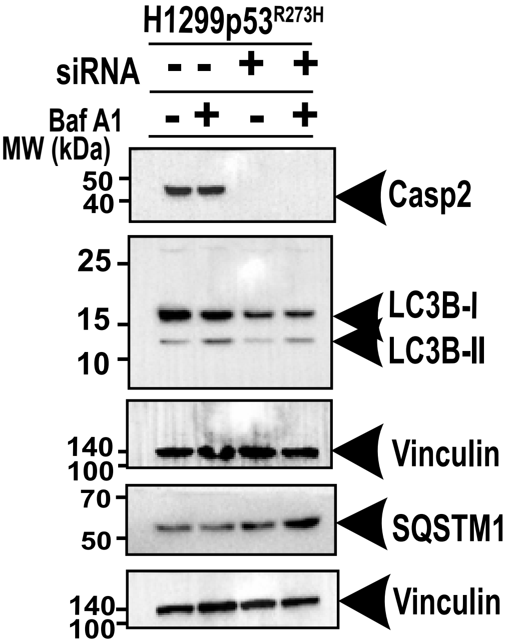
