## Supplementary Table 2 for "Caspase-2 protects against ferroptotic cell death"

```
#-----------------------------------------------------------------------------------------------
# MS Type | Cycle Id | Start IM [1/K0] | End IM [1/K0] | Start Mass [m/z] | End Mass [m/z]
#-----------------------------------------------------------------------------------------------
     MS1,      0,          - ,          - ,          - ,          - ,     -
   PASEF,      1,      0.9001,     1.2001,      800.00,      826.00,     -
   PASEF,      1,      0.6000,     0.9001,      400.00,      426.00,     -
   PASEF,      2,      0.9201,     1.2201,      825.00,      851.00,     -
   PASEF,      2,      0.6200,     0.9201,      425.00,      451.00,     -
   PASEF,      3,      0.9301,     1.2301,      850.00,      876.00,     -
   PASEF,      3,      0.6300,     0.9301,      450.00,      476.00,     -
   PASEF,      4,      0.9500,     1.2501,      875.00,      901.00,     -
   PASEF,      4,      0.6501,     0.9500,      475.00,      501.00,     -
   PASEF,      5,      0.9600,     1.2601,      900.00,      926.00,     -
   PASEF,      5,      0.6601,     0.9600,      500.00,      526.00,     -
   PASEF,      6,      0.9800,     1.2801,      925.00,      951.00,     -
   PASEF,      6,      0.6801,     0.9800,      525.00,      551.00,     -
   PASEF,      7,      0.9900,     1.2901,      950.00,      976.00,     -
   PASEF,      7,      0.6900,     0.9900,      550.00,      576.00,     -
   PASEF,      8,      1.0101,     1.3101,      975.00,     1001.00,     -
   PASEF,      8,      0.7100,     1.0101,      575.00,      601.00,     -
   PASEF,      9,      1.0201,     1.3201,     1000.00,     1026.01,     -
   PASEF,      9,      0.7200,     1.0201,      600.00,      626.00,     -
   PASEF,     10,      1.0401,     1.3401,     1025.01,     1051.01,     -
   PASEF,     10,      0.7400,     1.0401,      625.00,      651.00,     -
   PASEF,     11,      1.0601,     1.3601,     1050.01,     1076.01,     -
   PASEF,     11,      0.7601,     1.0601,      650.00,      676.00,     -
   PASEF,     12,      1.0701,     1.3701,     1075.01,     1101.01,     -
   PASEF,     12,      0.7701,     1.0701,      675.00,      701.00,     -
   PASEF,     13,      1.0901,     1.3901,     1100.01,     1126.01,     -
   PASEF,     13,      0.7901,     1.0901,      700.00,      726.00,     -
   PASEF,     14,      1.1001,     1.4001,     1125.01,     1151.01,     -
   PASEF,     14,      0.8001,     1.1001,      725.00,      751.00,     -
   PASEF,     15,      1.1201,     1.4201,     1150.01,     1176.01,     -
   PASEF,     15,      0.8200,     1.1201,      750.00,      776.00,     -
   PASEF,     16,      1.1300,     1.4301,     1175.01,     1201.01,     -
   PASEF,     16,      0.8300,     1.1300,      775.00,      801.00,     -
```

| CE [eV]
